# A stable cryogenic fluorescence microscope for correlative super-resolution light and electron microscopy

**DOI:** 10.64898/2026.04.18.719389

**Authors:** Soheil Mojiri, Joseph M. Dobbs, Ricardo Sánchez Loayza, Anna Kreshuk, Julia Mahamid, Jonas Ries

## Abstract

Cryogenic correlative light and electron microscopy (cryo-CLEM) enables visualization of biological specimens with molecular specificity while preserving near-native macromolecular structure. However, the severely limited resolution of conventional cryo-fluorescence microscopes restricts the accuracy of correlation with cryo-electron microscopy. Super-resolution cryogenic CLEM (SR-cryo-CLEM) offers a potential solution, but presents substantial technical challenges, including mechanical instability and ice contamination. Here, we introduce a modular cryogenic light microscope optimized for single-molecule localization microscopy (cryo-SMLM) that mitigates such limitations. The system is constructed primarily from off-the-shelf components, enabling straightforward and cost-effective assembly, and is operated using fully open-source Python software for flexible and customizable control. The mechanically and thermally stabilized architecture, combined with an axial focus-lock system, maintains sample positioning within a standard deviation of 40 nm. Ice contamination is minimized by imaging inside a purged enclosure, enabling prolonged acquisitions. Together, the platform provides robust localization precision, reproducible imaging performance, and an accessible solution for SR-cryo-CLEM.

## Introduction

The integration of fluorescence microscopy with electron microscopy for imaging of the same biological specimen under cryogenic conditions (cryogenic correlative light and electron microscopy, or cryo-CLEM) enables correlations of molecular specificity, from fluorescence imaging, with structural details of cellular architecture, from electron microscopy, while preserving samples in a vitrified, near-native state (Sartori et al. 2007; Schwartz et al. 2007; Lučić et al. 2013; Hampton et al. 2017; Wu et al. 2020). However, the three orders of magnitude gap in spatial resolution between these imaging modalities limits the accuracy and reliability of their correlation. Super-resolution cryo-CLEM (SR-cryo-CLEM) addresses this limitation by improving the resolution of the cryogenic fluorescence modality (Chang et al. 2014; Liu et al. 2015; Dahlberg and Moerner 2021).

Several cryogenic super-resolution techniques, including super-resolution optical fluctuation imaging (cryo-SOFI) (Moser et al. 2019), structured illumination microscopy (cryo-SIM) (Phillips et al. 2020; Li et al. 2023), and image scanning microscopy (cryo-ISM) (Wu et al. 2020), have demonstrated strong potential in cryo-CLEM applications. These approaches are particularly advantageous when rapid, low-dose imaging is required. However, most of these techniques provide only a two-to three-fold improvement in spatial resolution compared to diffraction-limited imaging, which remains insufficient for applications requiring nanometer-scale localization precision. Single-molecule localization microscopy (SMLM) overcomes this limitation by localizing individual fluorophores with high precision through the sequential activation of sparse subsets of emitters during long imaging sequences (Betzig et al. 2006; Rust et al. 2006). This approach can improve spatial resolution by one to two orders of magnitude compared to diffraction-limited imaging. Performing SMLM at cryogenic temperatures further enhances fluorophore photostability and reduces photobleaching, resulting in significantly higher photon yields and improved localization precision (Schwartz et al. 2007; Liu et al. 2015; Li et al. 2015, Weisenburger et al. 2017; Tuijtel et al. 2019; Dahlberg et al. 2018; Böning et al. 2020). However, performing robust and reproducible cryo-SMLM poses substantial technical challenges from the hardware perspective, with four key criteria being of particular relevance to SR-cryo-CLEM: first, the microscope must maintain high thermal stability and ensure constant cryogenic temperatures to maintain samples below the devitrification temperature (ca. 138K) (Dubochet et al. 1988). Second, it should provide minimal drift and vibrations over the long acquisition times required for cryo-SMLM (Wolff et al. 2016). Third, vitrified samples mounted on fragile EM grids must remain intact and free of ice contamination during transfer, imaging, and retrieval to ensure electron transparency in subsequent cryo-EM analysis (Dahlberg and Moerner 2021). Fourth, the system must provide flexible and user-friendly software for controlling excitation, photoactivation, and data acquisition.

Existing cryogenic widefield fluorescence microscopes can be categorized according to the sample environment: vacuum (closed) cryostats (Li et al. 2015; Hoffman et al. 2020; Böning et al. 2020; Hulleman et al. 2021), cryogenic immersion systems (Le Gros et al. 2009; Nahmani et al. 2017; Faoro et al. 2018; Faul et al. 2025), and cold nitrogen gas (open) cryostats (Tuijtel et al. 2019; Phillips et al. 2020; Dahlberg et al. 2020; Last et al. 2024; Schorb et al. 2017; Moser et al. 2019). Vacuum-based systems offer excellent mechanical stability, but require complex sample transfer workflows and demanding maintenance. Cryo-immersion systems enable imaging with higher numerical aperture objectives, and with an acceptable level of ice contamination during long image acquisition (Faul et al. 2025), but introduce additional challenges in sample handling in the immersion media, and can suffer from limited mechanical stability. These instabilities result from the short working distances typical of cryo-immersion objectives, which lead to strong thermal coupling between objectives and samples. Furthermore, the small margin between the cryostat operating temperature (base temperature) and the devitrification threshold in cryo-immersion systems limits the use of high excitation laser powers, whereas vacuum and open cryostats typically reach lower base temperatures and thus allow higher excitation powers (Mojiri et al., 2025).

Open cryostats offer simpler handling and easier sample access. However, they can suffer from accumulated ice contamination during long imaging sessions and often exhibit reduced mechanical stability (Wolff et al. 2016; Dahlberg and Moerner 2021). Mechanical instabilities here may arise from periodic drift associated with liquid-nitrogen refilling cycles, or from high-frequency vibrations transmitted by nitrogen boil-off or the LN_2_ injection systems (Dahlberg et al. 2018; Tuijtel et al. 2019; Phillips et al. 2020). These limitations are exacerbated by the long acquisition times required for SMLM, which typically involves recording thousands of sequential imaging frames to accumulate sufficient localizations for super-resolution reconstruction.

Additionally, cryogenic conditions also introduce fluorophore-specific challenges, including altered activation pathways and reduced switching efficiencies. These challenges restrict the palette of suitable labels and prolong the acquisition times (Tuijtel et al. 2019; Dahlberg et al. 2018; Last et al. 2025).

Beyond fluorophore photophysics, acquisition times are typically further prolonged because the excitation intensities that can be safely applied to samples plunge-frozen on conventional EM grids are highly limited (typically 20-300 W/cm^2^(Dahlberg et al. 2022; Last et al. 2023; Mojiri et al. 2025). These limits arise primarily from optical absorption and low thermal conductivity of common EM grid support films, which can lead to local heating of the frozen specimen and cause devitrification (Dahlberg et al. 2022; Last et al. 2023; Mojiri et al. 2025). Lower excitation intensities reduce fluorophore brightness and photoswitching rates (Tuijtel et al. 2019; Dahlberg et al. 2022), resulting in fewer detected photons per localization, and increasing the likelihood of overlapping point spread functions. Consequently, cryo-SMLM experiments typically require extended imaging sessions (tens of minutes to several hours) to accumulate sufficient localization events (Liu et al. 2015; Weisenburger et al. 2017; Tuijtel et al. 2019), further increasing susceptibility to sample drift and ice contamination.

Ice contamination in cryo-CLEM can originate from the accumulation of condensed water molecules during imaging, sample handling, or sample transfer between imaging modalities. Several strategies have been proposed to mitigate ice contamination in cryo-CLEM workflows. For example, purging of the cryogenic stage with heated nitrogen gas prior to imaging and sealing the system can significantly reduce ice accumulation (Arnold et al. 2016). Integrated light microscopy within a cryo-focused ion beam (cryo-FIB) platform, used for thinning samples to produce electron-transparent lamellae (Marko et al. 2007), or within transmission electron microscopy (TEM) systems, can reduce ice contamination by minimizing or eliminating sample transfers between modalities (Agronskaia et al., 2008; Faas et al., 2013; Gorelick et al. 2019; Li et al. 2023b; Yang et al. 2026). However, the confined geometry of these systems typically restricts the use of high-numerical-aperture objectives, thereby limiting fluorescence imaging resolution.

Despite substantial progress in recent years, integrating all these requirements into a cryo-SMLM platform that provides high thermal and mechanical stability, minimal ice contamination, reliable sample handling, and modular, extensible microscope control software remains technically challenging. Here, inspired by the design presented by Xu et al. 2018, we introduce a stable cryogenic super-resolution microscope with active temperature control of the objective, cryostat, and sample. Relative motion between the sample and objective is minimized by mounting the objective, sample, and translation stages within a cage-based framework. This configuration enables cryo-SMLM with minimal ice contamination during prolonged acquisitions. The microscope is built largely from off-the-shelf components, facilitating straightforward assembly. It also integrates a commercially available cryo-transfer shuttle for reliable handling of vitrified EM grids, and is operated using a newly-developed open-source modular software for instrument control and data acquisition. A fixed optical layout eliminates movement of the detection objective, ensuring a mechanically stable detection path. Compared to the system presented in Xu et al. (2018), our microscope offers several key advantages: (1) demonstrated reduction in contamination, along with successful SR-cryo-CLEM data acquisition; (2) fully open-source microscope control software; (3) a fixed optical layout that eliminates the need to move the objective during cryostat baking or sample transfer; (4) an integrated focus-lock system; (5) the use of a higher numerical aperture objective (NA = 0.9 versus 0.8); and (6) compatibility with a commercially available cryo-transfer shuttle, enabling reliable sample loading and unloading. The design, software, and operation manual are fully provided here.

## Methods and Materials

### Microscope setup

#### Sample and objective cage

We present our setup schematic in Fig. 1 and Fig. S1. We mounted our microscope objective, the translation stages, and the sample within a rigid cage assembly to minimize relative motion between components. This cage-like structure is built from Polyether ether ketone (PEEK) plates at its top and side walls, and a copper base plate at the bottom, where our translation stages and sample holder are mounted. The whole cage is connected to an aluminium plate mounted on the upper optical breadboard (600 mm × 600 mm, Throlabs) with an opening (100 mm × 100 mm). The aluminium plate is mounted using three location markers to allow for reliable optical alignment in case the cage structure is dismounted. Our long working distance upright objective (CFI TU Plan Apo EPI 100x, Nikon), is connected to the top PEEK plate using a copper adapter. The outer surface of the copper adapter is wrapped with heating foil and a temperature sensor (PT100) to maintain constant temperature of the objective (297.3 K) during the imaging experiments. In addition, to ensuring optical performance and facilitating mechanical stability, the objective temperature control prevents frost accumulation on the outer objective glass surfaces. The front part of the objective is covered with a plastic cap to thermally insulate the objective from the sample and cryostat. On top of the base plate, two translation cryo-stages (ANPx311, Attocube) enable lateral movement of the sample, followed by a one-dimensional lifting cryo z-stage (ANPz-102, Attocube) on top. The temperature of the sample holder (at 93 K) is monitored and controlled using a temperature controller module (model 335, LakeShore cryotronics) and a heat conductor device (ATC100/70, Attocube) consisting of two copper plates connected together with copper braids, a heater, and a temperature sensor (Fig. 2a). The heat-conducting plate containing the sensor and heater is attached on top of the z-stage. The other conducting plate is attached independently to the base plate to further improve conducting heat transfer between the sample and the cooling source.

**Figure 1.**
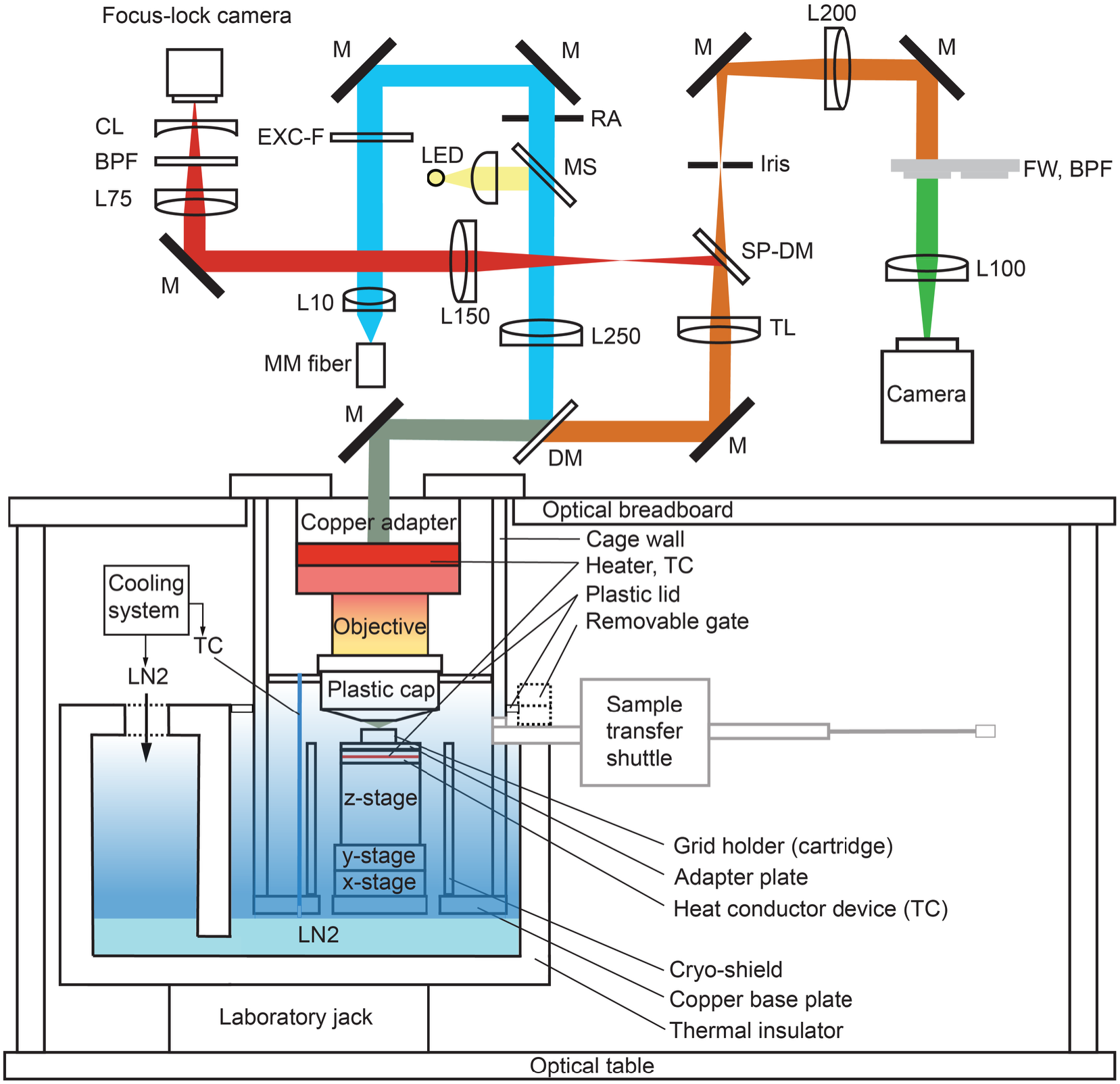
Schematics of the cryo-SMLM microscope. Top: wide-field fluorescence microscope. The excitation lasers shown in cyan output from a multi-mode (MM) optical fiber (150×150 µm, 0.39 NA, M103L05) are collimated using an aspheric lens (L10) and focused at the back focal plane of an objective using the focusing lens (L250). The fluorescence signal shown in brown is back-reflected through the objective, transmitted through the quad-band dichroic mirror (DM), and collected using the tube lens (TL). The fluorescence intermediate image created at the focal plane of the tube lens is projected on an sCMOS camera using a 4f telescope system comprised of the L200 and L100 lenses. The irises in the excitation and detection paths, respectively, adjust the size of the illumination and the detection field of view. The far-red fluorescence signal from dark-red fiducial markers, shown in red, is reflected to the focus-lock detection arm using a short-pass dichroic (SP-DM, cut-off at 697 nm). This signal is aberrated via a weak cylindrical lens (CL) and relayed to an auxiliary camera using another 4f telescope system made of lenses L150 and L75. Bottom: cryo-stage. The air objective is mounted in an upright configuration inside a cage system with a copper adapter, where heating foils and temperature sensors control its temperature. The cage, clamped on the upper optical table, is made of heat-insulating side walls and a conducting plate at the bottom, where the cryo-compatible translation stages and the sample are mounted. A liquid nitrogen (*LN*_2_) micro-dosing pump cools the stages and sample by injecting *LN*_2_ inside the thermal insulator, where the cage is immersed. The laboratory jack adjusts the height of the thermal insulator. The aluminium cryo-shield reduces the humidity around the stages and the sample. In addition, it enhances heat transfer by directing the cold nitrogen gas that enters from openings in the bottom base plate. The sample holder (cartridge) is transferred from the right side using the cryo-transfer shuttle and sits on a heat-conducting device containing a heater and a temperature sensor for controlling the sample temperature. L: lens (focal length specified in mm), M: mirror, RA: rectangular aperture, BPF: band-pass filter, EXC-F: excitation filter, FW: filter wheel, TC: temperature control.

**Figure 2.**
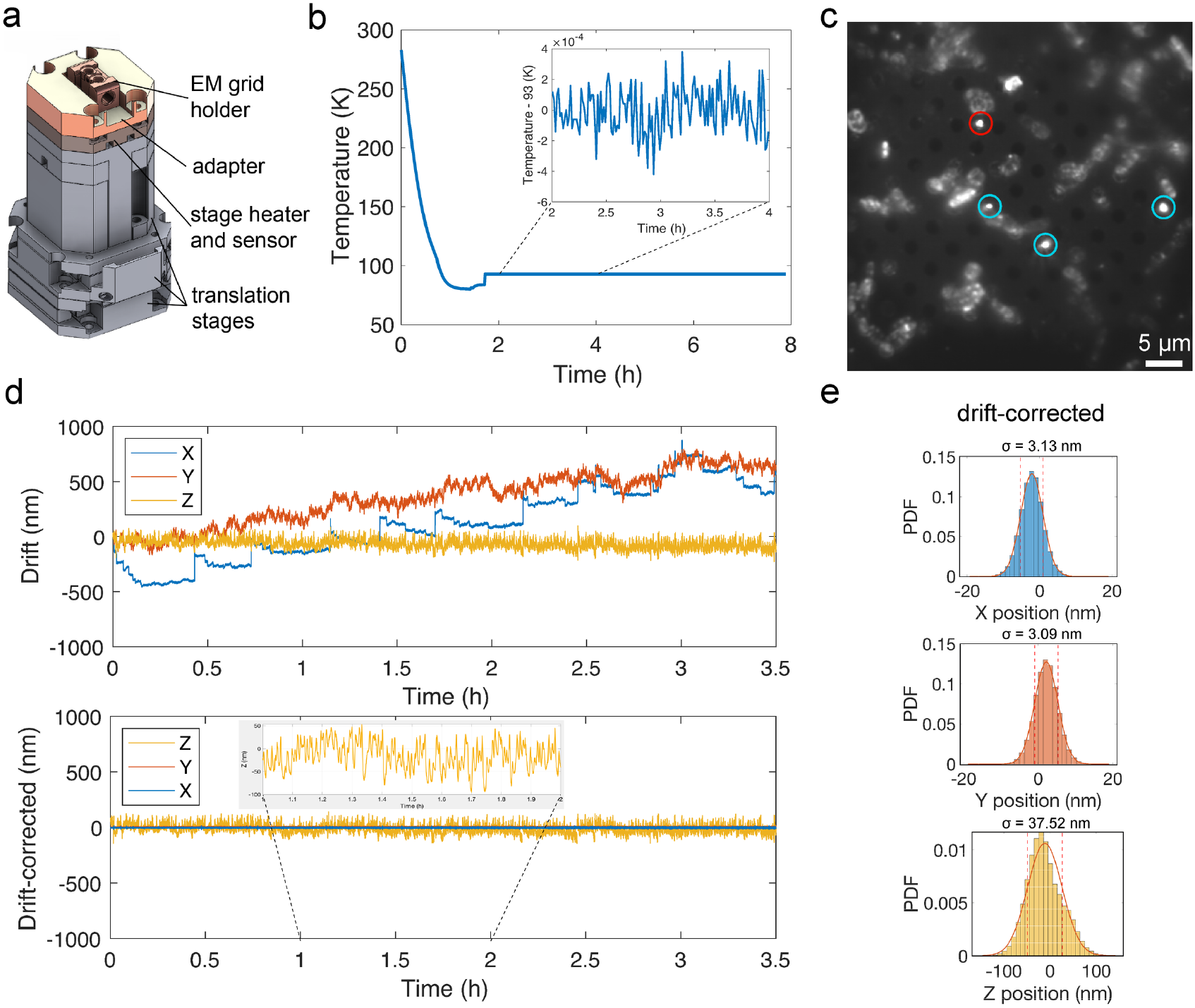
Thermal and mechanical stability of the cryo-SMLM open stat platform. a) From bottom to top: lateral and axial translation stages, the metallic plate containing the heater and temperature sensor, adapter for holding the sample cartridge, and cartridge with two slots to accommodate autogrid-mounted cryo-EM grids. b) Stage temperature cooling and thermal stability over more than six hours. The inset shows the average temperature values of every 50 binned data points over 2 hours. c) Vitrified E. coli cells on a cryo-EM grid together with tetra-spectral beads shown in red and cyan circles. d) Upper and lower panels showing correspondingly the average lateral drift and lateral drift-corrected positions (blue and red) measured for the beads encircled in c. Axial position values in yellow are the same in both panels, indicating the focus-lock precision. The inset in the lower panel plots the averaged axial position values of every 50 binned data points over 1 hour. e) Histograms and standard deviations of drift-corrected lateral positions and axial positions for the single bead highlighted in a red circle in c.

We used copper cartridges from Leica Microsystems as sample holders. These provided two slots with copper springs to hold autogrid-mounted EM grids. We incorporated two magnets at the bottom of the Leica cartridges to facilitate firm contact between the cartridge and the copper adapter plate, which is fixed on top of the heat-conducting plate.

#### Cryostat

The cage structure is surrounded by a heat-insulating container, into which liquid nitrogen is pumped using a *LN*_2_ micro-dosing system (model 915, Norhof). The *LN*_2_ micro-dosing system regulates the cooling power inside the cryostat by constant reading of the Norhof temperature sensor, which hangs freely inside the cryostat in the vicinity of the copper base plate. The pumping rate required to reach cooling temperatures below 95 K is mainly dependent on the size and insulation quality of the cryostat. It is critical to adjust the pump flow so that the *LN*_2_ level is kept just below the copper base plate to avoid transmitting high-frequency vibration from the boiling *LN*_2_ into the sample, and at the same time to provide maximum cooling power. The *LN*_2_ height measured from the bottom of our cryostat is nearly 12 mm. The *LN*_2_ pumping rate of our cooling system is set to 19 mbar. The gaps between the microscope objective and the cage walls, as well as between the cage walls and the cryostat, are covered with plastic lids to improve the thermal insulation of the cryostat and humidity control.

#### Humidity control

We perform cryo-SMLM in a laboratory equipped with a humidity control system that maintains the relative humidity in the range of 20-25%. However, this alone is not sufficient to prevent ice contamination during sample transfers and prolonged imaging sessions. Therefore, a plexiglass enclosure was built around the microscope to reduce humidity near the sample. Prior to cooling at 295 K, the enclosure (volume ∼ 0.2 m^3^) and cryostat were purged with nitrogen gas at an estimated flow rate of ≈ 100 L min^−1^ for nearly 40 min. Insulation of the cryostat from the environment inside the plexiglass, using plastic lids, further protects the sample from ice contamination. In addition, PTFE (Teflon) tubing was used to deliver nitrogen gas from the laboratory supply to the Plexiglas enclosure, as PTFE exhibits substantially lower permeability to water vapour than commonly used plastic pneumatic tubing such as PUN (polyurethane pneumatic) or PUR (polyurethane), thereby helping to maintain a dry purge environment.

#### Excitation and detection Optics

In the illumination part, we use an iBEAM SMART 488 (200 mW) and a Coherent Sapphire LPX 561 (500 mW) laser for excitation and an iBEAM SMART 405 (200 mW) for photo-activation.

In addition, a 638 nm diode laser (HL63193MG, Oclaro/Ushio) controlled by a laser engine box, as presented in Schröder et al. 2020, is added to the illumination path. The laser paths are overlaid on each other and coupled into a square multi-mode optical fiber (150×150 µm, 0.39 NA, M103L05) using long-pass dichroic mirrors (DMLP425, DMLP505, and DMLP605) and silver-coated mirrors. To achieve a homogeneous illumination across the field of view, speckles in the beam profile are significantly reduced by mechanically shaking the optical fiber attached to an elastic cord using a vibration motor (DC 1.5-6V, 16500 rpm, 22g, Sourcingmap) (Schröder et al. 2020). The fiber beam output, collected by a doublet lens, L10 (focal length 10 mm), is imaged by a doublet lens, L250 (focal length 250 mm) and the microscope objective, onto the sample plane, providing a homogeneous wide-field illumination.

Accurate placement and shaping of the excitation beam were essential for controlled illumination in our custom cryo-SMLM setup. To achieve this, a rectangular aperture (SP 60, Owis) was installed within a rotatable SM1-threaded lens tube (Thorlabs) and positioned at a field plane conjugated to the sample plane. In particular, this configuration enables the adjustment of both the dimensions and orientation of the illuminated region, allowing the excitation profile to be precisely aligned with the geometry and orientation of the sample during laser exposure. The filter (EXC-F) located between these two lenses is a four-line clean-up filter (390/482/563/640 HC Quad, AHF). The fluorescence signal collected by the objective and transmitted through a quad-band dichroic (4x dichroic mirror (F73-410, AHF)) is focused by a tube lens, TL (focal length 200 mm, TTL200-A, Thorlabs). The intermediate image plane of the tube lens is projected on a sCMOS camera (ORCA-Fusion BT, Hamamatsu) using a relay telescope 4f system comprised of Lenses L200 (focal length 200 mm) and L100 (focal length 100 mm). The effective camera pixel size is 136 nm. A motorized filter wheel (FW102C, Thorlabs) with six slots for positioning band-pass filters is positioned between the relay lenses, enabling switching between fluorescence channels.

#### Focus stabilization

Our EM grid samples (described below) included 200 nm tetra-spectral fluorescent micro-spheres, which we used for real-time correction of any slight mechanical drift along the axial dimension. The far-red fluorescence signal is partially split by a short-pass dichroic (SP-DM, HC 697 SP, AHF) and directed towards the focus-lock arm. The second 4f telescope system is comprised of Lenses L150 (focal length 150 mm) and L75 (focal length 75 mm), and transmits the tube lens image onto the auxiliary sCMOS camera (CM3-U3 50S5M, Chameleon3) with an effective pixel size of 133 nm. BPF in the focus lock arm indicates the far-red band-pass emission filter (700/50 nm, Semrock). We imposed astigmatic aberration on our far-red fluorescent PSF by exploitation of a weak cylindrical lens, CL (focal length -1000 mm, Thorlabs), to break the symmetry of the PSF above and below the focus. A z-stack of the astigmatic PSF is collected to attribute each aberrated PSF image to a certain defocused axial value (details in the supplementary file). The focus-lock routines are implemented in a Python-based program that analyzes the PSF and controls the stage accordingly (Fig. S9).

In detail, focus-locking functions by analyzing a small cropped region of interest (ROI; 32–80 px) from the auxiliary camera in real time. For each incoming frame, the intensity is projected along the x- and y-axes, and the resulting 1D profiles are fitted with a Gaussian function plus a linear background. The Gaussian standard deviation *σ*_*y*_ and *σ*_*y*_ are extracted, and their ratio r=*σ*_*x*_/*σ*_*y*_ is used as the focus metric, with values near unity indicating optimal focus. A scalar Kalman filter is applied to the raw ratio values to suppress noise and reject outliers (Kalman 1960), yielding a stable estimate of the focus metric. The controller continuously monitors the filtered ratio, and deviations from the optimal value trigger corrective Z-steps of the stage in the appropriate direction, restoring the sample to the focal plane (details in the supplementary file). Lateral drifts are corrected in post-processing using the same bead images.

### Microscope control and data acquisition software

We implemented a novel, fully Python-based microscope control system using a custom hardware abstraction layer and PyQt5 GUI, following the design principles of the Python-Microscope (“Microscope”) library (Susano Pinto et al. 2021). This approach provides a flexible and modular framework that allows direct control of cameras, stages, lasers, and filter wheels, while enabling user-friendly interfaces for live imaging, multi-step Z sweeps, and automated navigation. Our graphical interface, built using PyQt5, allows users to manage acquisition parameters, visualize live camera streams, and interactively define navigation points (Fig. S2). Acquisition tasks are executed asynchronously to maintain responsive control during automated experiments. The software leverages the filter-wheel and laser modules of the Python-Microscope library for hardware abstraction (Susano Pinto et al. 2021). The software supports translation of the x, y, and z stages individually (Fig. S4) and Z-axis sweeps for PSF analysis and volumetric imaging (Fig. S5), multi-position navigation (Fig. S10), and synchronized image acquisition from two cameras (Fig. S3, S9). Peripheral devices, including lasers (Fig. S6, S7) and filter wheels (Fig. S8), are controlled through dedicated modules, enabling flexible experimental configurations. The microscope control software, including the Python packages, is available in (Github, https://github.com/kreshuklab/CryoSResCLEMcontrol). A detailed description of our software GUI is provided in the Supplementary file.

### Microscope preparation and data acquisition

The humidity inside the plexiglass environmental enclosure was first reduced to below the sensitivity of our hygrometer (0.1%) by purging the enclosure with nitrogen gas. During this drying period of nearly 40 min, and prior to cooling, the objective temperature controller was switched on and set to a few degrees above room temperature (typically 298 K) to prevent condensation during the experiments; the objective was allowed to reach thermal equilibrium. The XY stages were then centered by locating the middle of a dummy EM grid loaded on the Leica cartridge at room temperature, and the cryo Z-stage was lowered to its bottom-most position to provide clearance for cartridge loading.

The cryogenic stages and sample holder were cooled to approximately 80 K over ∼ 75 min with LN_2_ flow at 21 mbar (adjustable by Norhof micro-dosing cooling system). After thermal stabilization, a vitrified EM grid, pre-clipped into an autogrid cartridge, was mounted in the precooled, modified Leica cartridge and inserted into the microscope using the Leica cryo-transfer shuttle. Low-intensity white-light illumination was switched on for visual guidance, and the cryo Z-stage was raised until the grid surface came into focus. As the sample approaches the objective, its temperature increases slightly. After focusing, the LN_2_ flow pressure was reduced to 19 mbar, and the stage temperature was set to 93 K to enable active thermal feedback control during imaging. The system was allowed to re-equilibrate for 10 min before data acquisition.

The center of the inserted grid was identified and registered in the Navigator GUI, and ROIs were located and marked. Fluorescence imaging parameters, including excitation and activation laser intensities and pulsing, camera exposure times, and the appropriate band-pass emission filter, were then adjusted for acquisition. After imaging, the cryo Z-stage was lowered to its bottom-most position, and the XY stages were recentered. The cartridge was subsequently unloaded, and the imaged grid was retrieved and stored for downstream cryo-EM analysis.

### Sample preparation

An IPTG-inducible FtsZ-rsEGFP2 plasmid with chloramphenicol selection was constructed using Gibson assembly from a plasmid expressing FtsZ-mEos2 with a T7 promoter (Addgene #49764, gift from Jie Xiao (Buss et al. 2013)) and a plasmid expressing rsEGFP2 (Addgene #102879, gift from Stefan Jakobs (Grotjohann et al. 2012)). *E. coli* BL21 DE3 cells were transformed with this plasmid and grown to 600 nm optical density (OD_600_) 0.6 in M9 minimal medium, at 37° C, with 20 *μ*g/ml chloramphenicol and 300 rotations per minute shaking. IPTG was added to 0.1 mM, and cells expressed the fusion construct for 15 minutes. 3.5 *μ*l of the suspension was applied to UltrAufoil R2/2 200 mesh gold grids (Quantifoil Micro Tools) which had been plasma cleaned (Fischione M1070) for 1 minute with a 75:25 argon: oxygen gas mix. 4 *μ*l of sample, including a 1:10 dilution of 200 nm tetraspectral beads (TetraSpeck™ Microspheres, 0.2 *μ*m, fluorescent blue/green/orange/dark red, Thermo Fisher Scientific) was applied to the grids, and these were vitrified in an ethane/propane cryogen after blotting from the back (37 C, 40% humidity, 1.3 seconds) in a Leica GP1 plunger (Leica Microsystems).

### Cryo-EM data acquisition and processing

Cryo-EM and cryo-electron tomography (cryo-ET) were performed using a Titan Krios G4 operated at 300 kV, with a Falcon 4i detector, Selectris energy filter, and a 100 *μ*m objective aperture. Data were collected using SerialEM (Mastronarde 2003). Single projections (Fig. 3b) were acquired at 6.28 angstroms per pixel (Å/px, nominal magnification 19,500x) and a fluence of 30 e^-^/Å^2^. Square maps (Fig. 4a,d,e) were acquired at 26.39 Å/px (nominal magnification 4,800x). Tilt series (Fig. 4f) were acquired from -60 to +60 degrees at 3° increments, using the dose-symmetric tilt scheme (Hagen et al. 2017), a fluence of 3.1 e^-^/Å^2^ per tilt, and at 3.74 Å/px (nominal magnification 33,000x). Tomograms were preprocessed in WarpTools (Tegunov et al. 2026), aligned with Aretomo2 (Zheng et al. 2022), and reconstructed in WarpTools at 24 Å/px.

**Figure 3.**
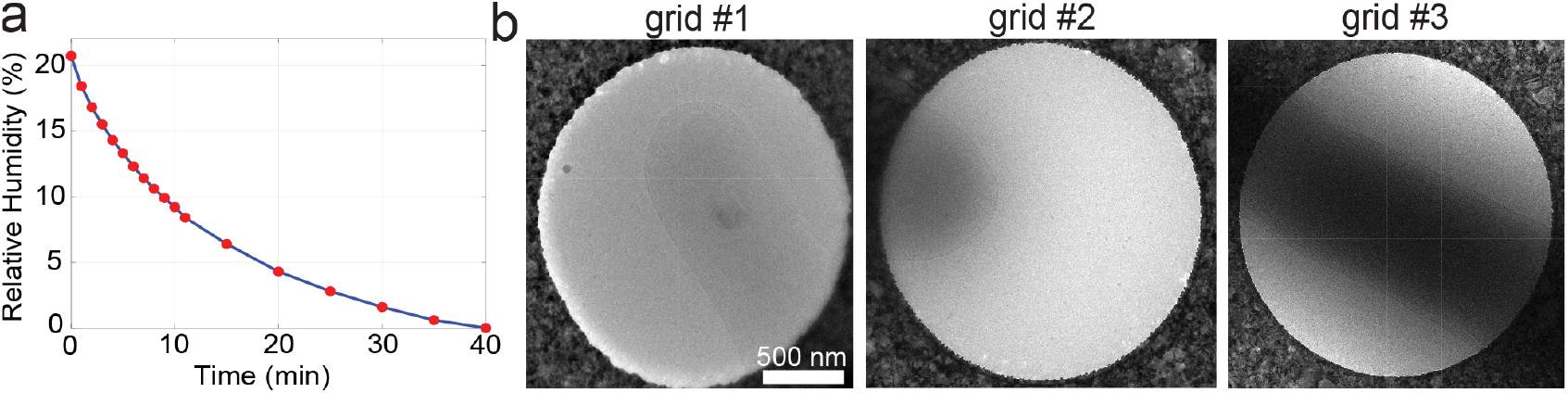
Reduced humidity and ice contamination in the cryo-SMLM open stat platform. a) Relative humidity measured inside the plexibox. b) Cryo-EM micrograph from a random hole in grid #1 after loading, incubation for 5 minutes, and unloading from the cryo-SMLM microscope. Two additional random holes from two different grids (#2 and #3), are shown after being imaged independently in the cryo-SMLM microscope for 3.5 hours.

**Figure 4.**
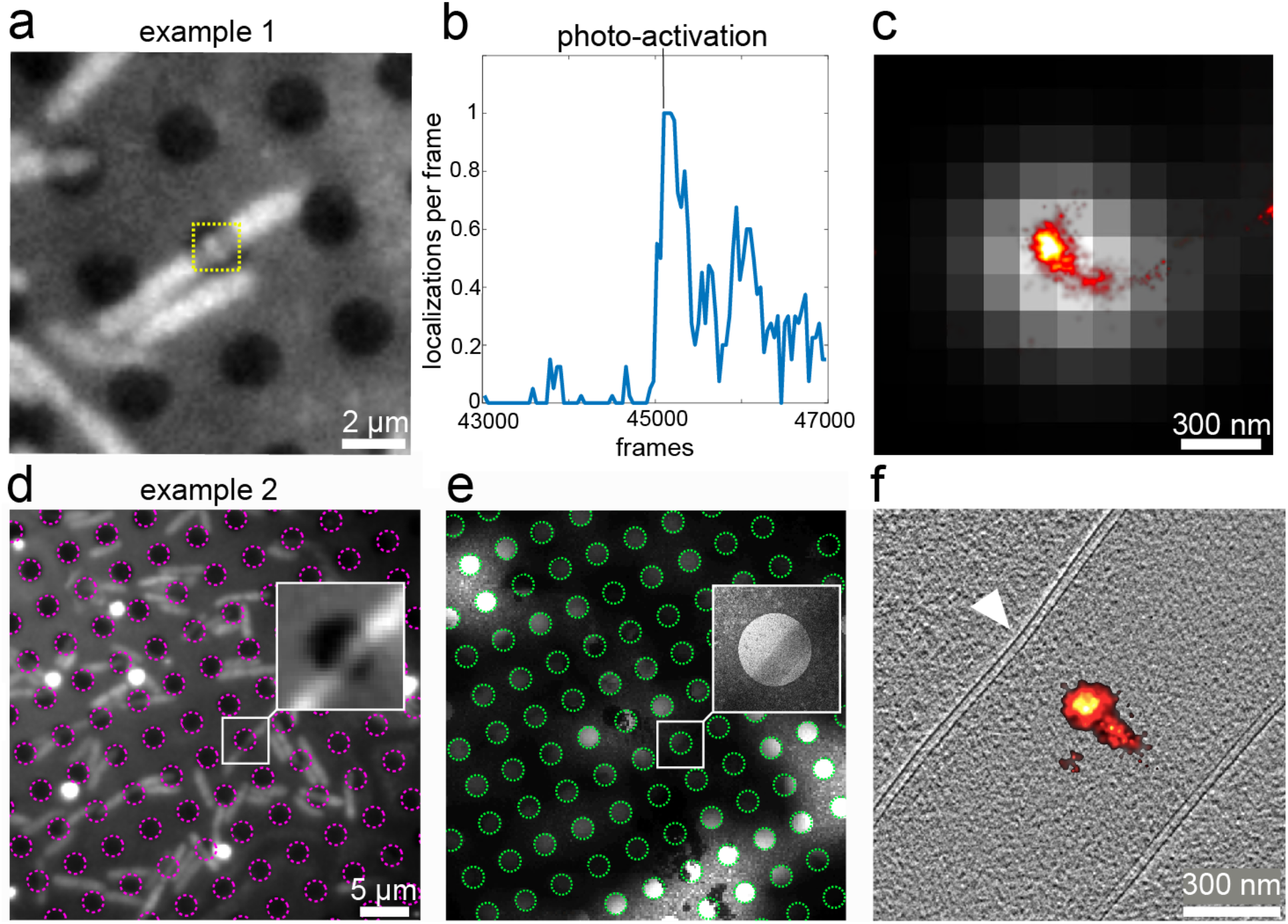
SR-cryo-CLEM results. a) First frame of the fluorescence imaging sequence showing several E. coli cells on an Ultrafoil grid. b) Increase in the number of detected localizations following photoactivation (at the indicated time point). c) Diffraction-limited rendering of FtsZ–rsEGFP2 (gray) and the corresponding SMLM reconstruction (red-hot color scale) from the ROI indicated by the yellow dashed square in (a). d) Cryo-fluorescence image of another batch of cells, with grid holes highlighted by magenta dashed circles. The inset shows a zoomed-in view of a single cell. e) Low-magnification cryo-EM image and inset of the same regions as in (d), with detected grid holes indicated by green dashed circles for correlation with the fluorescence image. f) Central slice from the cryo-ET volume corresponding to the white square in (e) (grayscale), correlated with the cryo-SMLM image of FtsZ–rsEGFP2 (red-hot color scale). The white triangle indicates the cell constriction site.

## Results

### Microscope Thermal and Mechanical Stability

The results of the thermal and mechanical stability measurement of our microscope are presented in Fig. 2. Fig.2 a shows the position of the temperature sensor with respect to the Leica cartridge and translation stages. Fig. 2b illustrates the temperature measurements over a period of 8 hours and demonstrates the temperature stability of the plate containing the stage heater and the temperature sensor below the sample holder, after the initial cooling period of approximately 75 minutes (*LN*_2_ pump flow set to 22 mbar reach the stage temperature of 80 K). The stage temperature was then set to 93 K. The flow pressure was set to 19 mbar, so sufficient cooling power is provided, and the liquid nitrogen level is kept just below the bottom base plate. The inset (Fig. 2b) presents the temperature value averaged over a bin of 50 values for a time window of two hours, showing that the standard deviation of the reference 93 K temperatures is ±0.0015 *K*.

To assess the mechanical stability of our microscope, we imaged TetraSpeck fluorescent beads added to an *E. coli* suspension and plunge frozen on an EM grid (Material and Methods), shown in Fig. 2c. The upper panel of Fig. 2d represents the average measured drift of four beads shown in the circles in Fig. 2c in the three spatial dimensions, imaged over 3.5 hours with temporal sampling of 100 ms. The axial positions were actively stabilized during image acquisition. Small-amplitude (∼ 250 nm), periodic displacements observed along the X direction are attributed to nitrogen pump operation. These drifts were highly reproducible and did not impact data quality, as they were efficiently removed during post-processing.

The inset shows the averaged axial positions binned over 120 values for 1 hr, representing the precision of our active focus-lock system. Lateral drift was corrected using the SMAP plugin implemented in MATLAB (Ries et al., 2020). Specifically, the ‘Drift correction Beads’ plugin was applied to fiducial markers to estimate the temporal x–y drift trajectory, which was subsequently smoothed with a sliding-window averaging filter and subtracted from all localization coordinates. The lateral drift correction algorithm can be accessed at: (Github, https://github.com/jries/SMAP/tree/master/plugins/%2BProcess/%2BDrift).

The two upper panels of Figure 2e illustrate the spatial distribution of bead localizations after drift correction (corresponding to the red circle highlighted in Fig. 2c).

Over the 3.5-hour acquisition period, we obtained a lateral standard deviation of ∼ 3 nm and an axial standard deviation of ∼ 40 nm. This mechanical stability allows us to consistently perform long acquisitions that are typically required for SMLM.

### Minimal ice contamination

By purging the microscope plexiglass enclosure with nitrogen, the relative humidity inside the imaging environment decreased from 20% to below 0.1% within approximately 40 min (Fig. 3a), allowing us to perform measurements over multiple hours with minimal ice contamination. To assess ice contamination arising solely from loading and unloading plunge-frozen grids into the cryostat, a grid was inserted into the imaging position, kept inside for 5 minutes, and then imaged by cryo-EM (Fig. 3b, #1). The results indicate minimal ice contamination during these handling steps. To evaluate longer-term effects, a few grid holes from five random squares of two additional grids were imaged after extended residence in the cryostat. Representative images show consistently low levels of ice contamination (Fig. 3b), with grid #2 and grid #3 maintained in the imaging position for approximately 3.5 hours.

### SR-cryo-CLEM

We evaluated the performance of our microscope using SMLM of the FtsZ protein in *E. coli* cells plunge-frozen on Ultrafoil EM grids. FtsZ forms the Z-ring that constricts bacterial cells to effectuate cell division (Barrows and Goley 2021), and was exogenously expressed as an FtsZ-rsEGFP2 construct. The autofluorescence background was first photobleached for 20 minutes with 488 nm illumination at an intensity of 20 W/cm^2^. Imaging was then performed using a 488 nm laser at 93 W/cm^2^, which is close to the devitrification threshold previously determined for this grid type at a base temperature of 93 K (Mojiri et al. 2025). Photoactivation of rsEGFP2 was achieved using continuous-wave 405 nm illumination at 1.8 W/cm^2^. Single-molecule localization and subsequent analysis were performed using SMAP (Ries et al., 2020). A region of interest containing several beads visible throughout the reconstructed dataset was first selected, and drift correction was applied as described above. Single-molecule localizations were identified using SMAP’s default peak detection algorithm and subsequently fitted with a 2D Gaussian model using standard fitting and filtering parameters. Localization precision was estimated within SMAP based on photon counts and background noise. Signals detected in consecutive frames were merged when separated by less than 35 nm and dark for a maximum of 1 frame.

For visualization, localizations were rendered using a red-hot lookup table, applying a localization precision filter of 0–30 nm. Poorly fitted localizations were excluded based on a threshold applied to the normalized log-likelihood. The final SMLM renderings were exported as TIFF files for correlation with low-magnification electron microscopy images. Fig. 4a–c shows an example of cryo-SMLM imaging of an E. coli cell without electron microscopy (EM) correlation: a representative fluorescence image of non-bleached FtsZ–rsEGFP2 within a grid hole is shown in Fig. 4a. The labeled FtsZ–rsEGFP2 signal is visible as a line inside the grid hole within the yellow dashed square. Upon illumination with 405 nm light starting at frame 45,000, a marked increase in detected localizations within the ROI (yellow dashed square in Fig. 4a) is observed (Fig. 4b). This burst in localization events confirms that the observed blinking originates from rsEGFP2. From the total of 51,000 acquired frames (example 1), the SMLM reconstruction shown in Fig. 4c was generated using the last 6,000 frames (100 ms exposure time), during which photoactivation enriched for specific blinking events and improved contrast of localized single molecules. The resulting SMLM image (red-hot color scale in Fig. 4c) exhibits an average localization precision of approximately 30 nm. For comparison, the corresponding diffraction-limited rendering is shown in gray in the background.

Figure 4d presents another cryo-SMLM example of FtsZ–rsEGFP2 in an E. coli cell, where 45,000 frames were recorded, and photoactivation was initiated at frame 12,000. The SMLM reconstruction shown in Fig. 4f was generated from the last 15,000 frames, during which the background signal was low and only photoactivated frames were included. The fluorescence and electron microscopy datasets were correlated using the grid hole pattern (Anderson et al. 2018) (magenta and green dashed circles in Fig. 4d and Fig. 4e, respectively). The insets in Fig. 4d and Fig. 4e show magnified views of the same grid hole containing the cell imaged by light microscopy and EM. The resulting correlated SMLM image of FtsZ from the highlighted ROI (white squares) is shown in Fig. 4f together with a central slice from the corresponding high-magnification cryo-ET volume of the E. coli cell. Together, these results demonstrate that our microscope enables high-precision SR-cryo-CLEM.

## Discussion

Here, we presented a modular and mechanically stable cryogenic super-resolution microscope, designed to improve the reproducibility and accessibility of cryo-SMLM for SR-cryo-CLEM. Our mechanical design minimized vibrations and drift during long acquisitions, enabling robust focus stabilization through active correction of axial drift using a user-selected fiducial bead. Using a single fiducial bead for focus locking was sufficient to achieve stable axial correction during extended SMLM acquisitions on plunge-frozen EM grids. In addition, these fiducial markers were used for accurate post-acquisition lateral drift correction. In addition, we developed a modular Python-based control platform that enabled flexible operation of the microscope. The software can be further extended with capabilities such as automated grid tiling, intensity modulation of excitation and activation lasers, and integration of application-specific hardware. For compatibility with downstream EM analysis, we systematically optimized key experimental parameters affecting contamination in the cryogenic imaging environment and evaluated ice quality using cryo-EM. Our optimized setup provided minimal ice contamination, allowing extended imaging sessions without compromising sample quality for correlative cryo-EM experiments.

Despite these improvements in the microscope setup, several limitations remain. The use of an air objective restricted the achievable numerical aperture (NA=0.9) compared with cryogenic immersion systems (NA=1.15-1.2). While our configuration simplified the mechanical decoupling and contributed to the observed stability, higher-NA optics in cryo-immersion microscopy workflows could further improve photon collection efficiency and cryo-SMLM throughput (Nahmani et al. 2017; Faoro et al. 2018; Faul et al. 2025). In addition, excitation intensities are also constrained by grid heating (∼0.02–0.3 kW/cm^2^), which may induce devitrification and limit performance for dim or inefficiently switching fluorophores, remaining well below typical room-temperature SMLM conditions (1–500 kW/cm^2^, Diekmann et al. 2020). A number of recent studies suggest that optimized grid materials can substantially increase the tolerable excitation intensity. For example, custom silver-coated grids (Dahlberg et al. 2022) and gold support films (illuminated at 640 nm) tolerate significantly higher excitation powers than commonly used carbon or gold films illuminated at shorter wavelengths (Mojiri et al. 2025). In particular, gold films illuminated at 640 nm can withstand intensities nearly twenty times higher than carbon films at the same wavelength. Because vitreous ice absorbs less light than support films, grids with larger hole fractions and adaptive illumination strategies that confine excitation to grid holes may further reduce heating.

Cryo-SMLM is also highly relevant for workflows involving cryo-FIB–milled samples, where thinning biological material into electron-transparent lamellae (100–300 nm) enables high-resolution cryo-ET (Marko et al. 2007). Recently, it was shown that exploiting an interferometric optical signal within the ice layer, specifically its depth-dependent modulation, can improve targeting accuracy during cryo-FIB milling (Sica et al. 2026). In a similar manner, cryo-SMLM has the potential to enhance targeting precision during milling, while subsequent cryo-SMLM on the resulting thin lamellae could further aid in selecting regions of interest and precisely localizing molecules for cryo-ET data acquisition (as explored in a forthcoming application study (Mojiri et al., manuscript in preparation).

However, the typical inclination of lamella (8–15° relative to the grid plane, Schaffer et al. 2017) introduces a height gradient that could exceed the optical depth of focus for high-NA objectives. Future extensions of the present microscope could address this limitation by incorporating a rotatable cryogenic stage to align lamellae perpendicular to the optical axis, or by implementing a tilted detection pathway, similar to that used in single-objective selective plane illumination microscopy (SO-SPIM (Galland et al. 2015)), to match the imaging plane to the lamella inclination.

It is important to note that the intensity of excitation also influences the photophysics of fluorophores and the efficiency of SMLM acquisition (Diekmann et al. 2020). Higher laser powers increase fluorophore brightness and photoswitching rates, reducing acquisition duration and thereby further benefiting stability and contamination control. Developing optimized imaging protocols that balance excitation intensity, fluorophore switching behavior, and devitrification limits will therefore be important for robust and high-throughput cryo-SMLM. Our current microscope configuration also supports multi-color cryo-SMLM through sequential imaging of spectrally-distinct fluorophores. Simultaneous dual-color imaging could be implemented by modifying the detection path to spectrally separate emission signals using appropriate dichroic mirrors and beam splitters (Gregor et al. 2021; Power et al. 2024). Parallel acquisition would shorten the total imaging times, reduce ice contamination, and further enhance stability during extended measurements.

In summary, we described a modular cryogenic super-resolution microscope for cryo-SMLM imaging with markedly reduced drift and ice contamination during extended acquisitions. By combining active temperature control, robust axial drift correction, optimized sample handling, and open-source control software, the system provides an accessible platform for SR-cryo-CLEM workflows. Continued advances in fluorophores, grid materials, illumination strategies, and automated correlation methods are expected to further improve imaging speed, robustness, and throughput for nanoscale studies of vitrified biological specimens.

## Acknowledgments

This work was supported by the Chan Zuckerberg Initiative (CZI) Visual Proteomics Program (Grant No. 2021-234620) and the EMBL. R. S. has received funding from the European Union’s Horizon 2020 research and innovation programme under the Marie Skłodowska-Curie grant agreement No. 945405. The authors acknowledge the facilities provided by the EMBL Cryo-EM Platform and the EMBL Imaging Centre. We thank the group members of J.M. and J.R. for their insightful discussions and valuable input throughout this work.

## Supplementary information

### 1 Setup Layout

**Figure S1.**
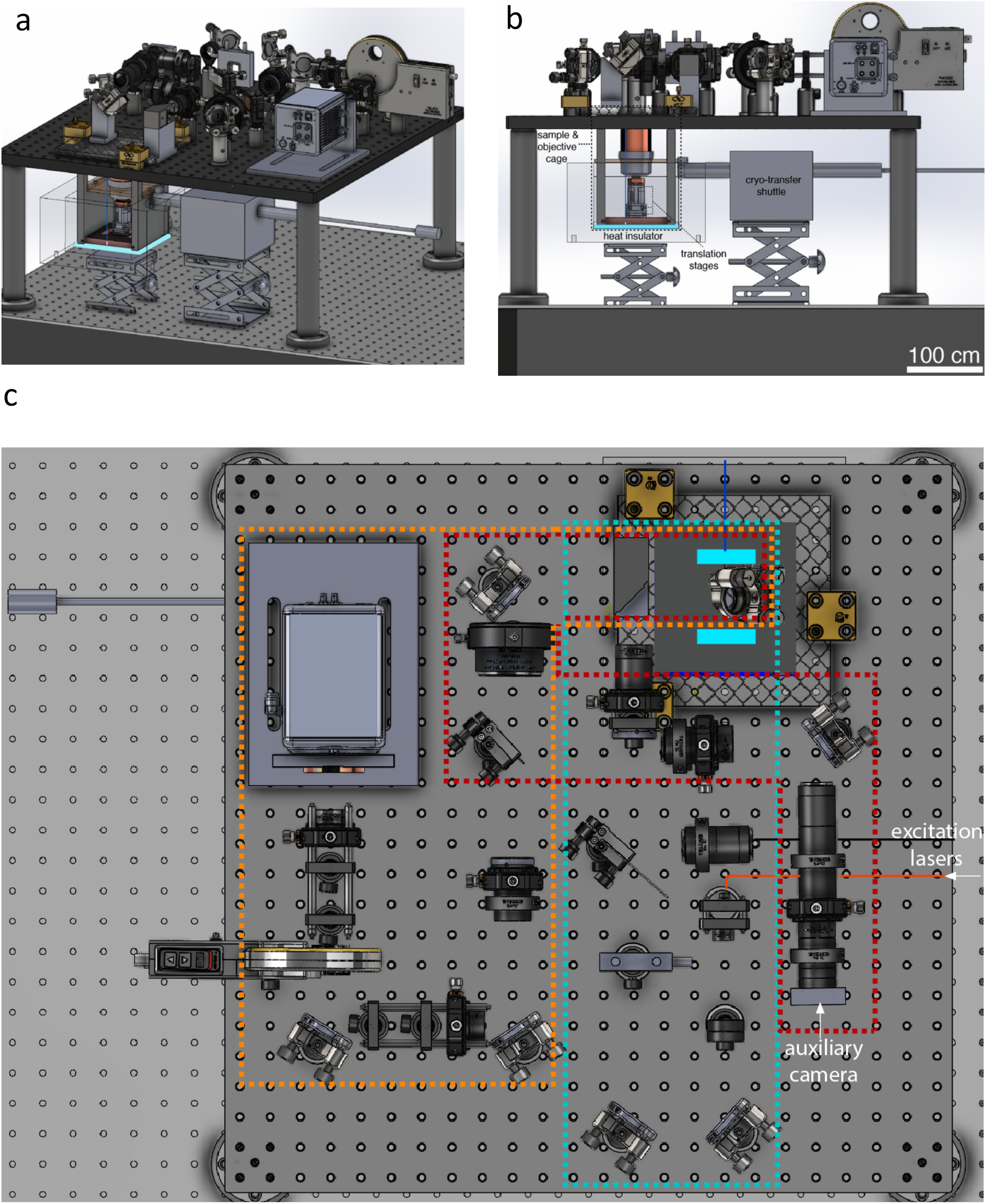
Microscope layout based on SolidWorks files. a) angled, b) side, and c) top views of the layout, correspondingly. The cyan, orange, and red dashed lines in the top-view panel indicate the illumination, detection, and focus-lock paths, respectively.

### 2 Microscope control software

#### 2.1 Software Implementation

The software was designed with a focus on both performance and usability. It employs multiple threads to acquire, process, and visualize data concurrently.

The software is written in Python 3.10, which is the latest version supported by our camera API/SDK, and uses PyQt for the graphical user interface and multithreading infrastructure, including thread management, inter-process communication via signal/slot mechanisms, and data sharing. Hardware communication relies on several Python libraries: *python-microscope* (Pinto et al. 2021) and *microfpga* (Deschamps et al. 2023) for optical components; *hidapi* and the proprietary *Spinnaker* SDK for camera control; and *pyserial* for serial communication with the motorized stages. Data processing is performed using standard scientific Python libraries, including *NumPy* (Harris et al. 2020), *SciPy* (Virtanen et al. 2020), *Numba* (to accelerate computationally intensive tasks) (Lam et al. 2015), *scikit-image* (Van der Walt et al. 2014), and *scikit-learn*. The software supports the NDTIFF format through the *ndstorage* Python library, and metadata are stored in plain CSV files. Development was carried out using the Spyder integrated development environment.

The software architecture is event-driven. Acquisition and processing tasks are executed in background worker threads, typically one per hardware component or processing task, while the main GUI thread remains responsive. Signals and slots are used to manage start and stop commands, handle task completion callbacks, and propagate updates to camera and stage states.

#### 2.2 Key Functionalities

- Dual camera image acquisition with live preview, record, and file management.
- Coarse and fine Z-stack acquisition.
- Multi-position stage navigation and automated movement.
- Focus tracking and automated compensation.
- Laser switch and intensity control, filter wheel control.
- Asynchronous worker-based acquisition to maintain responsive GUI operation.
- Export of experimental parameters and navigation positions for reproducibility.

#### 2.3 Microscope control and GUI

**Figure S2.**
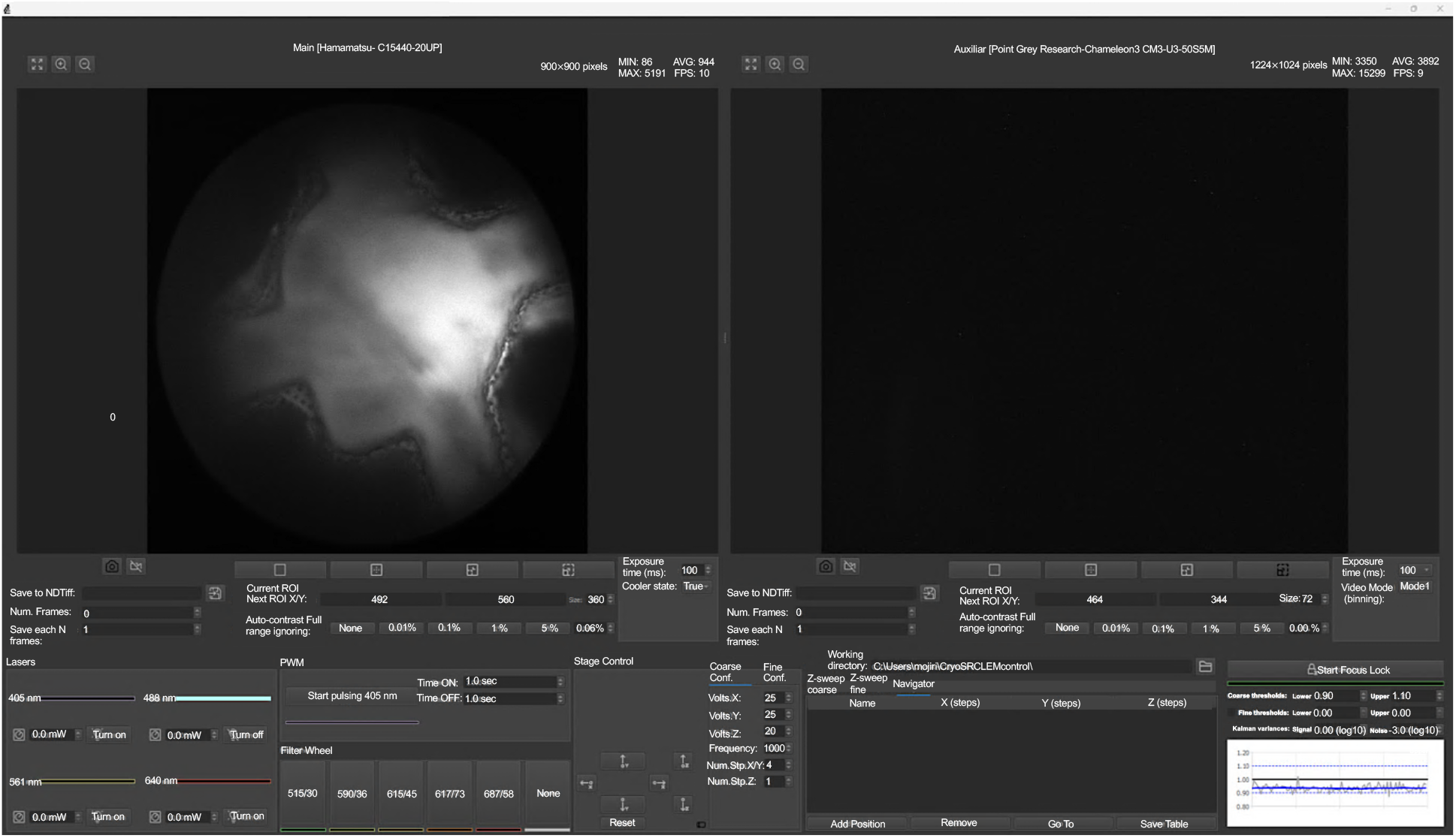
Overview of the graphical user interface of the control software (controls detailed below).

##### Camera Control

The software supports simultaneous control of two cameras (“main” on the left of Fig. S2 and “auxiliary” on the right). The user can start or stop live acquisition and frame recording for each camera independently. Filename management is integrated, allowing image saving from either or both cameras during frame recording and automated z-sweeps. Camera frames are saved in NDTiff format. Fig. S3 shows the annotated GUI controls for selecting and adjusting the camera ROI. A brief tooltip description appears when hovering over each icon. The numbered items correspond to:

1. View full image: fits the full sensor image to the display.
2. Zoom in.
3. Zoom out.
4. The pixel at the top-left corner of the camera sensor, defined as the origin (x = 1, y = 1)
5. Acquire a snapshot
6. Enable/disable live imaging mode
7. Show ROI: displays the currently selected ROI.
8. Pick ROI center: after selecting this tool, hold the Ctrl key and scroll the mouse wheel forward to enlarge the ROI (scroll backward to reduce its size). Once the desired ROI size and position are set, press button 6.
9. ROI in: confirms and activates the selected ROI.
10. ROI out: deselects the ROI and returns to the full-chip view.

**Figure S3.**
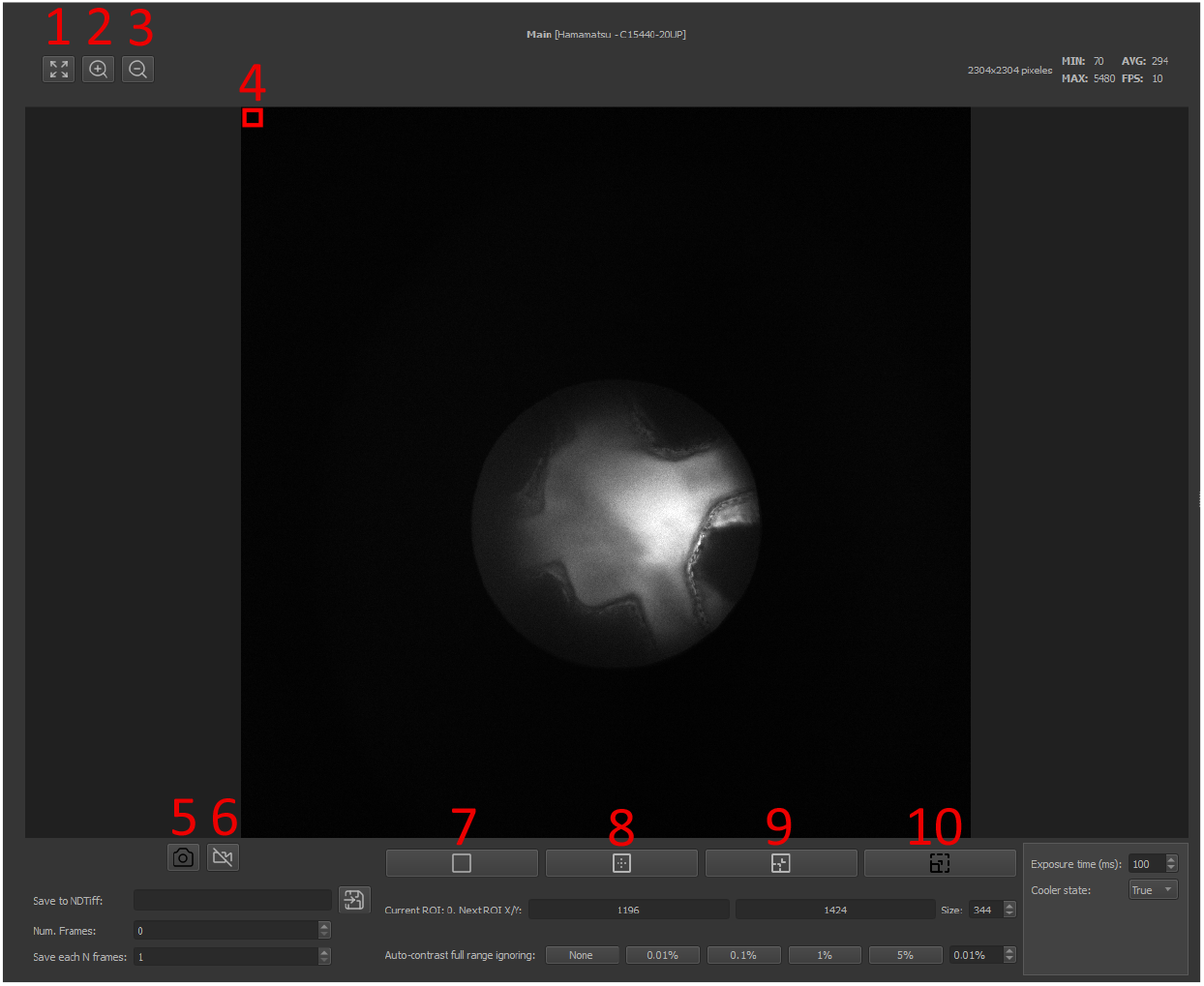
The main camera control. Numbers shown in red indicate the ROI selection and zoom-in/out options. Numbers on the top right represent the shown ROI size, minimum (MIN), maximum (MAX), and average (AVG) pixel values, and frames per second (FPS). The black camera icon enables acquiring a single frame and the white camera icon enables and disables live imaging mode. ‘save to NDTiff’ enables allocating a folder name for frame saving. The number of saved frames can be set by ‘Num. Frames’. Time-lapse image saving is possible using ‘save each N frames’, where the default value of ‘N=1’ translates to saving all frames sequentially. The center position of the selected ROI, and its size, can be correspondingly set using ‘Next ROI X/Y’ and ‘size’. ‘Auto-contrast full range ignoring’ allows for adjusting the image contrast based on pixel values and signal-to-noise ratio. Besides the pre-defined values of ‘None’, 0.01%, 0.1%, 1%, and 5%, a manual contrast value can be set in percent in the same row. ‘Exposure time (ms)’ can set the camera exposure time. ‘Cooler state’: ‘True’ enables water cooling, while ‘False’ enables internal fan (air) cooling.

##### Control of translation stages

Sample positioning is done by the translation stage movement in two different modes. The first mode involves coarse movement (step mode), utilizing discrete step commands with a specific step size (voltage) in either the forward or backward direction. The step size is dependent on different parameters, including the load on the stage, temperature, and stepping frequency.

In our setup, the minimum voltage required to produce visible sample movement at cryogenic temperatures (78–100 K) is approximately 20 V for the x stage and 15 V for the y and z stages. In step mode, the minimum, maximum, and incremental voltages are set to 15 V, 150 V, and 1 V, respectively. At 93 K, the measured step sizes are about 200 nm for the x stage at 20 V and 150 nm for the y stage at 15 V.

Fine movement (offset mode) uses an offset voltage within the (0-150) V range. The smallest offset voltage can be set to 0.1 V. As the voltage-distance calibration is dependent on the sample load and temperature, it should be independently conducted for each hardware setup. The maximum offset voltage step in the program is set to 5 V. We calibrated the voltage–distance relationship of our z-stage by imaging fluorescent beads located on the top and bottom surfaces of a coverslip with a nominal thickness of 170 μm (diameter: 2.5 cm, refractive index: 1.52). Because imaging was performed with an air objective, the axial displacement of the focal plane differs from the mechanical stage displacement due to the refractive index mismatch between air and glass. Accounting for this effect, the effective mechanical travel corresponding to the coverslip thickness is reduced by a factor of approximately (1/1.52). Based on this correction, we estimate a stage step size of ∼15 nm per 0.2 V offset (offset mode) when moving upwards at 93 K. The same measurement at room temperature with step voltage (step mode) with a voltage of 15 V (upwards) results in a ∼ 100 nm step.

**Figure S4.**
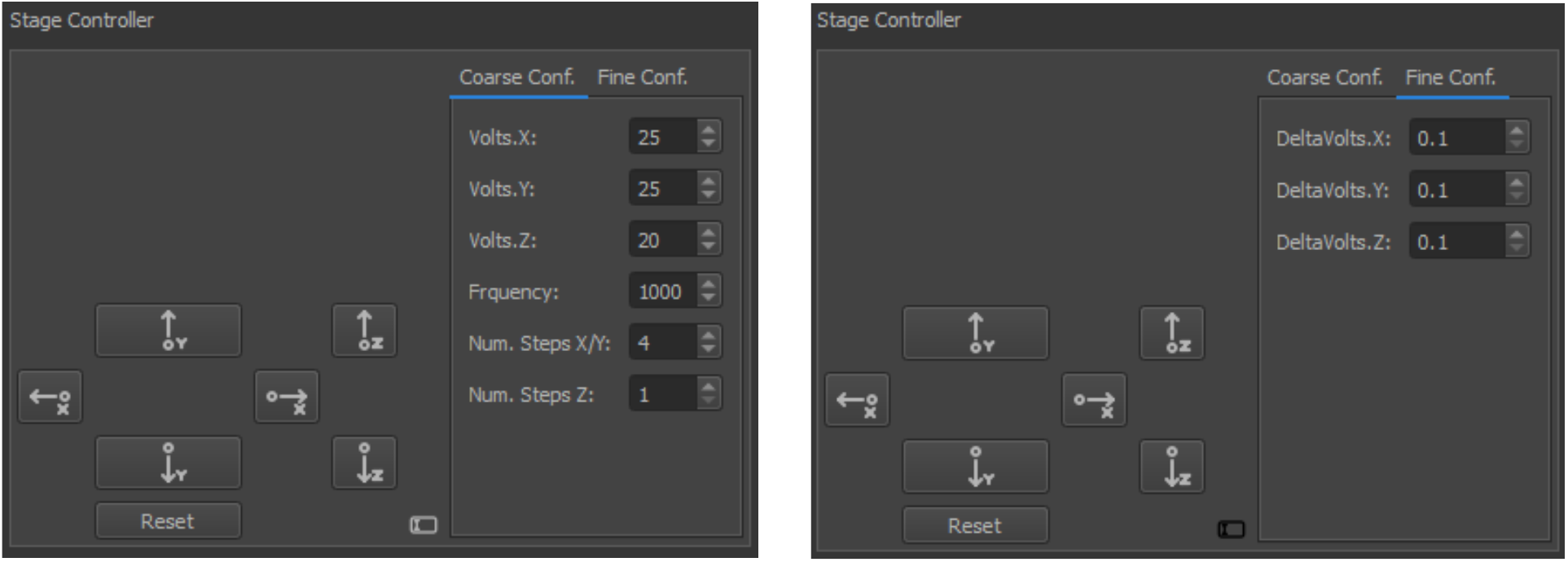
Translation stage control. The coarse movement (step mode, left panel) and fine movement (offset mode, right panel) controls. Step sizes can be adjusted by setting the applied voltages (in volts) to each axis. Frequency (in Hz) determines the speed at which the stepping is performed. One can determine the number of steps per click on the software arrows on the left by setting ‘Num. Steps X/Y’ for the lateral movement and ‘Num. Steps Z’ for the axial movement. Stepping can also be easily controlled using the computer keyboard, via up and down arrows (y-axis), left and right arrows (x-axis), or page up and page down keys (z-axis). Fine stepping can be done by applying offset voltages in the tab named ‘Fine conf.’, where the step size can be determined by setting the offset voltages (DeltaVolts.X,..) in Volts. An offset voltage of 0 V or 150 V represents the lower and upper limits for the stage driver. When either limit is reached, the system stops stepping automatically according to the programmed ‘stop stepping’ function. The ‘Reset’ button switches the stages off and on again, in case the offset voltage reaches 0 or 150 V, accidentally.

##### Z-axis control (z-sweep)

Coarse (step mode) z-sweep: Allows large-step relative movements or scans along the z-axis to find the focal plane of the imaging or to obtain and evaluate the experimental PSF of the microscope. The user can specify the number of steps, voltage increments (equivalent to the step size), and delays between movements. The sign of the given voltage in ‘step size’ determines the direction of movement (plus sets scanning from bottom to top, and minus sets scanning from top to bottom).

Fine (offset mode) z-sweep: The same functionality as in coarse z-sweep, but with small-step, high-resolution movements for precise z-scanning. Both sweep types support image saving from one or both cameras.

**Figure S5.**
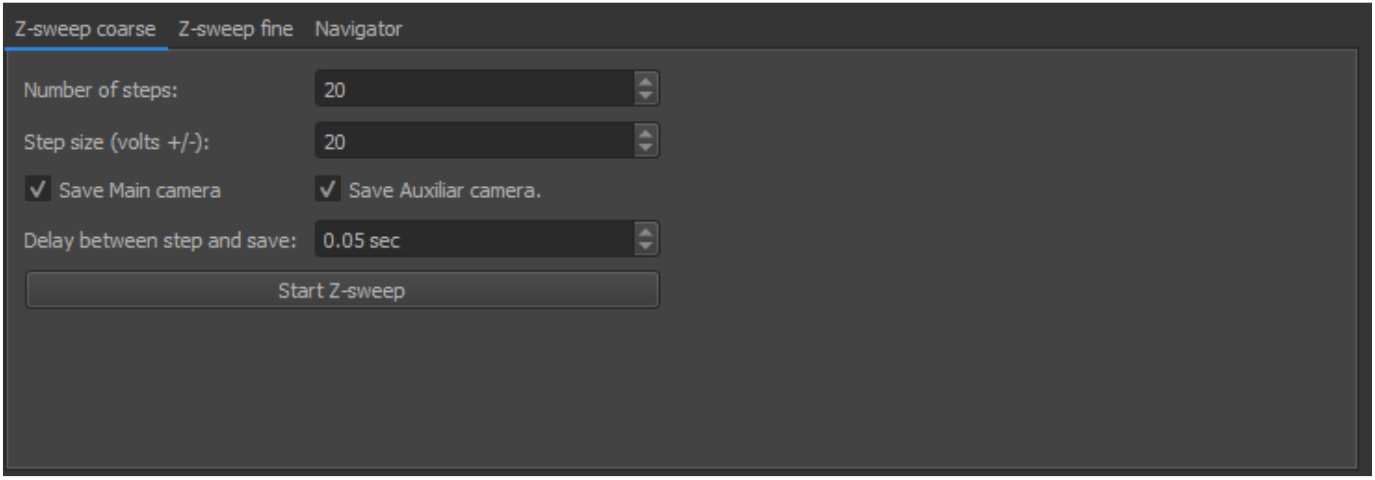
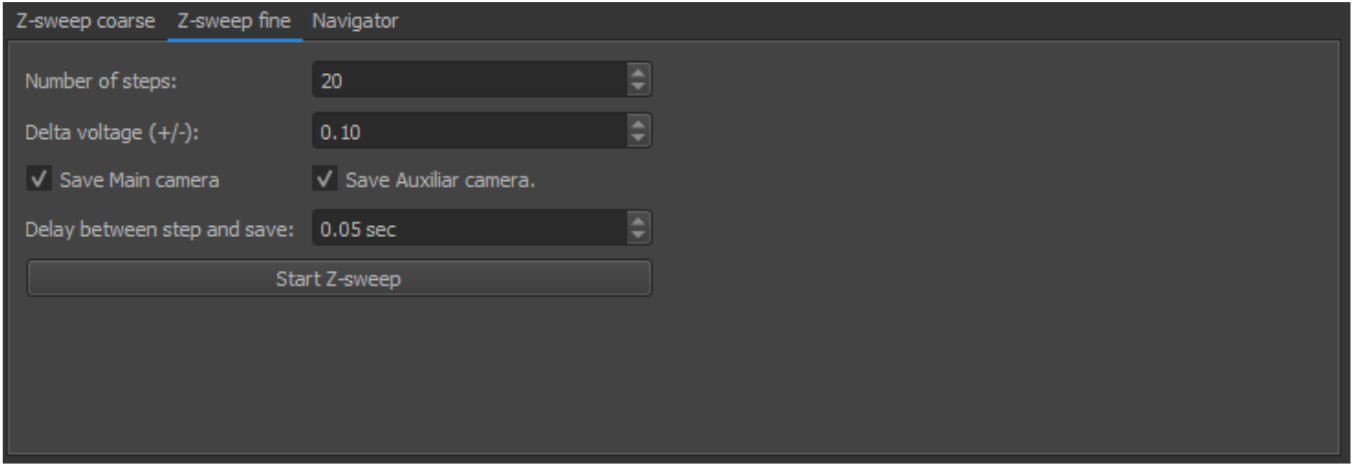
Upper and lower panels show coarse and fine z-sweep control tabs, respectively.

##### Laser illumination control

Individual control of multiple excitation laser sources (405, 488, 561, 640 nm) through dedicated widgets with pulse-width modulation for precise control of laser intensities, enabling precise laser dosing/excitation. For each nominal laser power setting defined in the software, the actual laser intensity at the sample should be determined by measuring the delivered power at the objective’s front focal plane.

**Figure S6.**
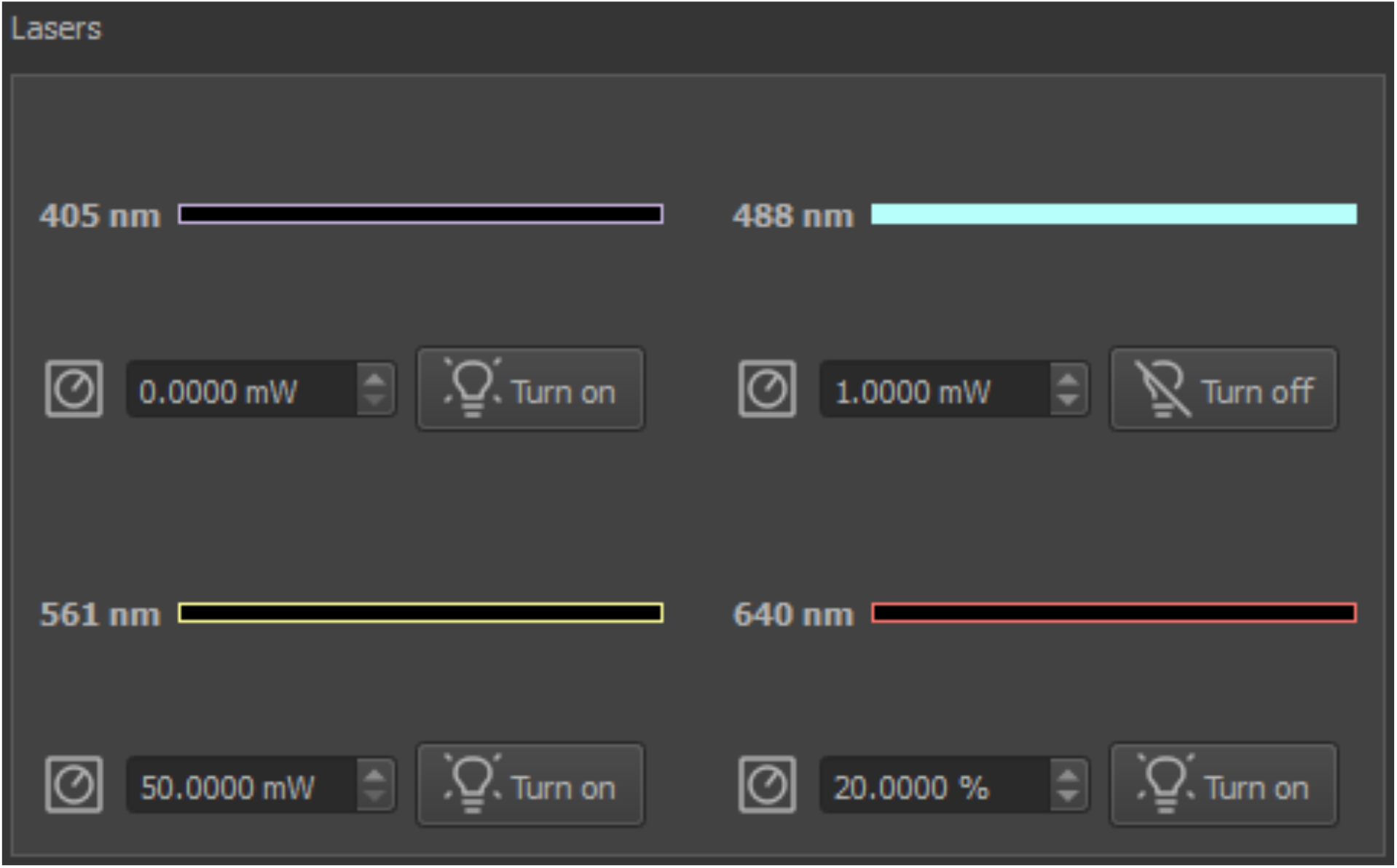
Laser switch and intensity control of: 405 nm and 488 nm iBeam Smart Toptica lasers with 200 mW power, 561 nm Coherent Sapphire LPX with 500 mW power, and 640 nm diode laser with a power mapped in a 0-100 % range.

##### Pulsing 405 nm laser illumination

The 405 nm laser can be operated in either continuous-wave (CW) or pulsed mode to enable controlled photo-activation in cryo-SMLM experiments. Pulsed operation provides precise temporal control over activation, which is particularly useful for managing activation density in samples with high autofluorescence background.

**Figure S7.**
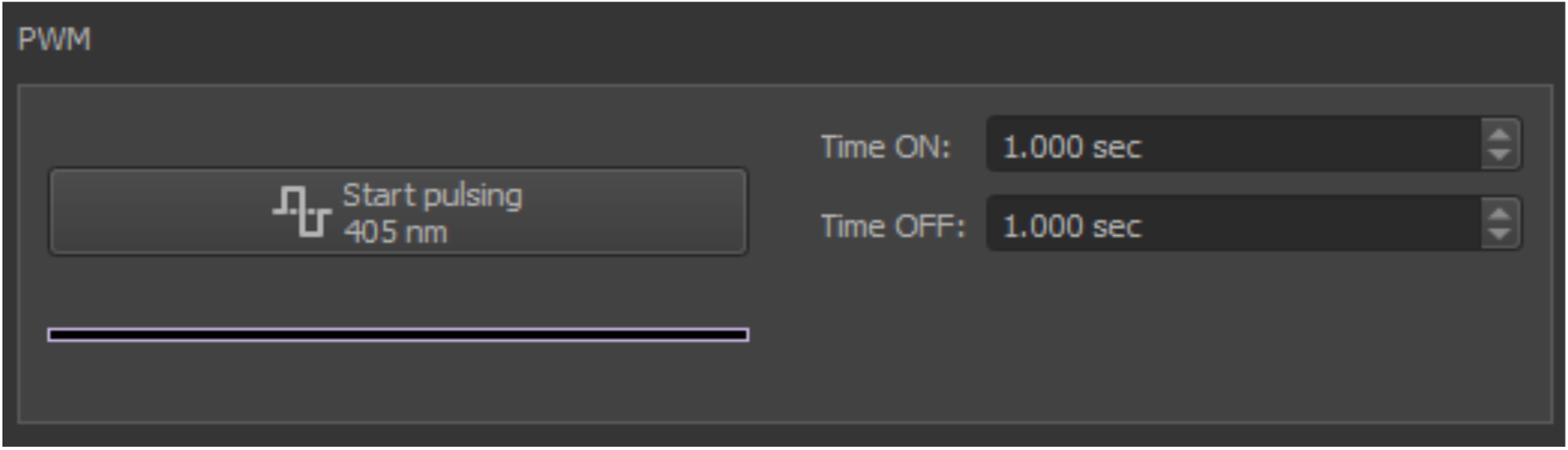
Pulsing mode for the 405 nm laser, specific for modular photo-activation. The pulse duration and repetition rate can be set using the Time ON and Time OFF options. The shortest ‘ON’ time (pulse duration) can be set to 1 ms.

##### Emission filter wheel

Software-controlled filter wheel selection with visual feedback.

**Figure S8.**
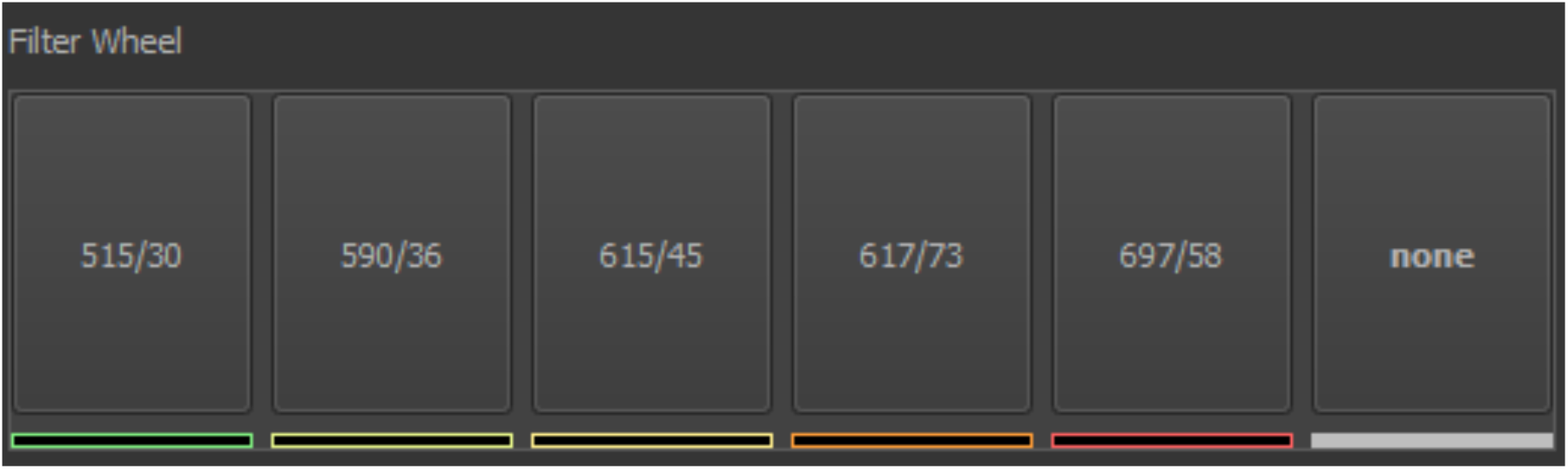
Emission filter wheel selection tab for five different emission band-path filters.

##### Focus-lock activation and real-time monitor

The focus-lock system maintains the Z-focus by tracking the astigmatic point-spread function (PSF) of a fiducial feature (typically a fluorescent bead; see Material ad Methods). The system continuously measures the x- and y-axis widths (standard deviations) of the bead image. When their ratio deviates from the calibrated in-focus value, the stage is adjusted accordingly. It is thus important that the feature is spherical.

The ROI selected on the auxiliary camera should contain a single bead and typically be 32–80 pixels in size. When beads are densely distributed, the minimum ROI size of 32 × 32 pixels can be used. This size ensures that a bead remains inside the ROI even under lateral drift of up to 1 µm (∼8 pixels) over a period of ∼4 hours, while still allowing reliable astigmatic PSF fitting along both axes for focus-locking. The full sensor sizes are 2304 × 2304 pixels for the main camera and 2448 × 2048 pixels for the auxiliary camera.

Because beads can lie at different heights with respect to the support film, the chosen fiducial bead should be located near the main imaging ROI viewed on the primary camera. This ensures that maintaining focus on the bead also maintains focus on the sample. Fig. S8 shows the GUI controls for selecting and adjusting the ROI on the auxiliary camera and focus-lock settings.To reduce jitter from noisy measurements, a Kalman filter is applied to the ratio, producing a smoothed focus estimate used for stage corrections. The GUI provides two adjustable parameters under “Kalman Variances”:

Signal: expected variability of the true focus ratio. Higher values make the system respond faster to real focus changes; lower values make it smoother but slower. Numbers are set in log 10 base.

Noise: expected measurement uncertainty. Higher values reduce stage jitter by ignoring noise; lower values make the system react more strongly to each frame, possibly causing jitter. Numbers are set in log 10 base.

Adjust these parameters to balance responsiveness and stability: increase Signal for faster corrections or Noise for smoother operation. Typical starting values are small Signal (≈5×10^−4^) and moderate Noise (≈1.0).

**Figure S9.**
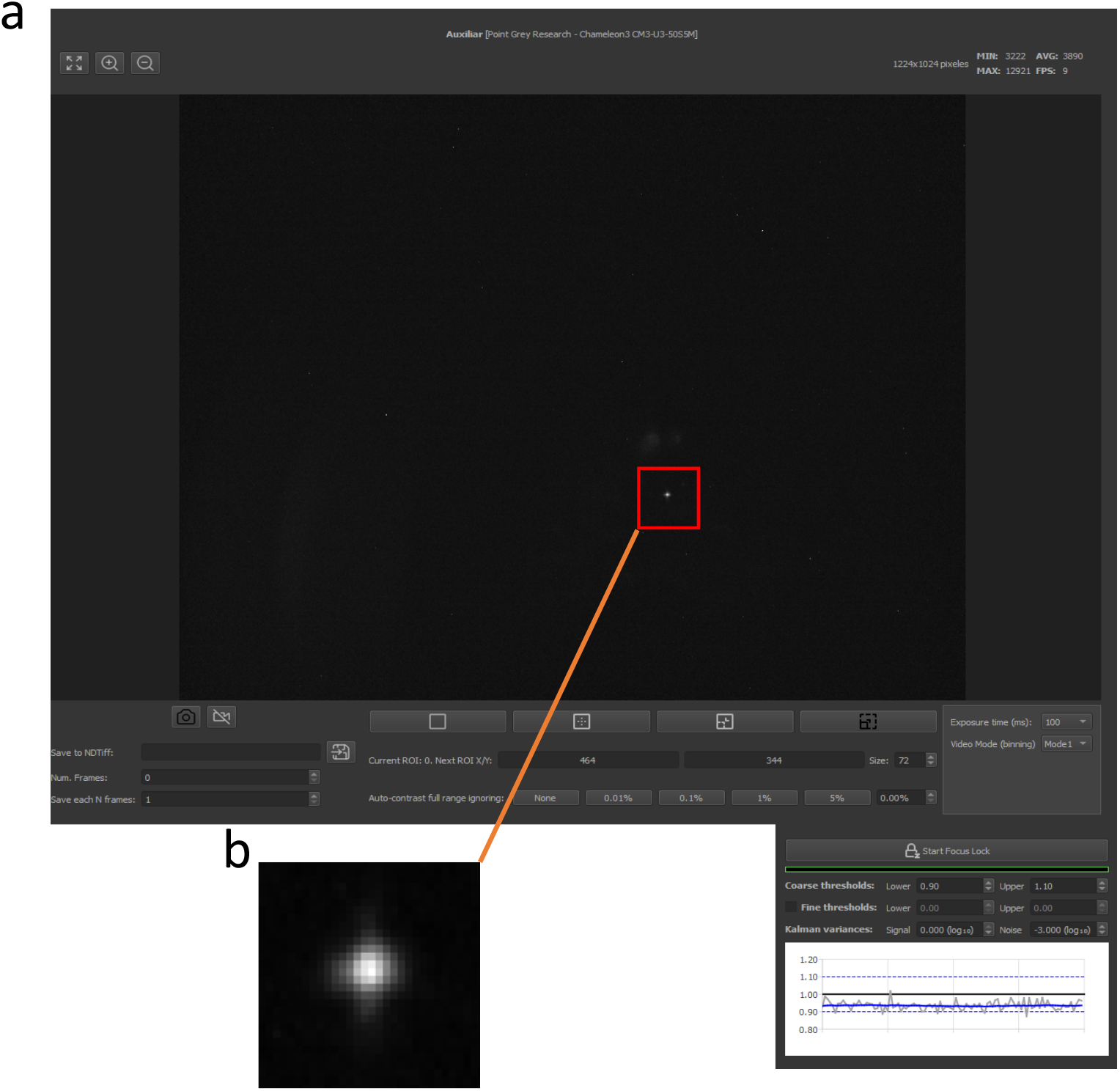
a) Auxiliary camera and focus-lock settings. All similar buttons to those in Fig. S3 do the same functionalities as described for the main camera, except the cooling option for the auxiliary camera is not provided. In addition, only five different exposure times and two pixel binning modes can be set using the specific camera model used as the auxiliary camera in this work, with a default exposure time set to 100 ms and camera video binning mode set to ‘Mode0’: no Binning, ‘Mode1’: binning 2 pixels. ‘Start Focus Lock’ and ‘Stop Focus Lock’ enable and disable focus lock accordingly. Focus-lock can be conducted in coarse or fine stepping mode. In each mode, the upper and lower bounds of out-of-focus ratio, *σ_x_*/*σ_y_*, can be set accordingly. In Kalman variances, one can preset ‘signal’ and ‘noise’ depending on the image signal-to-noise ratio for a more robust fitting. When Focus-lock is activated, the current out-of-focus ratio is shown in the lower panel, where the gray plot shows the actual measured value, and the blue line shows the Kalman-filtered value. The blues dashed lines represent the upper and lower bounds of axial extent, where the focus is locked within. b) Enlarged image from the single bead image with astigmatic PSF at focus.

##### Stage Navigation

The software provides a multi-position navigation table for storing and revisiting positions in x, y, and z directions. Users can add, remove, or rename positions, and automatically move the stage to any selected entry. Positions can be saved to CSV for reproducibility.

**Figure S10.**
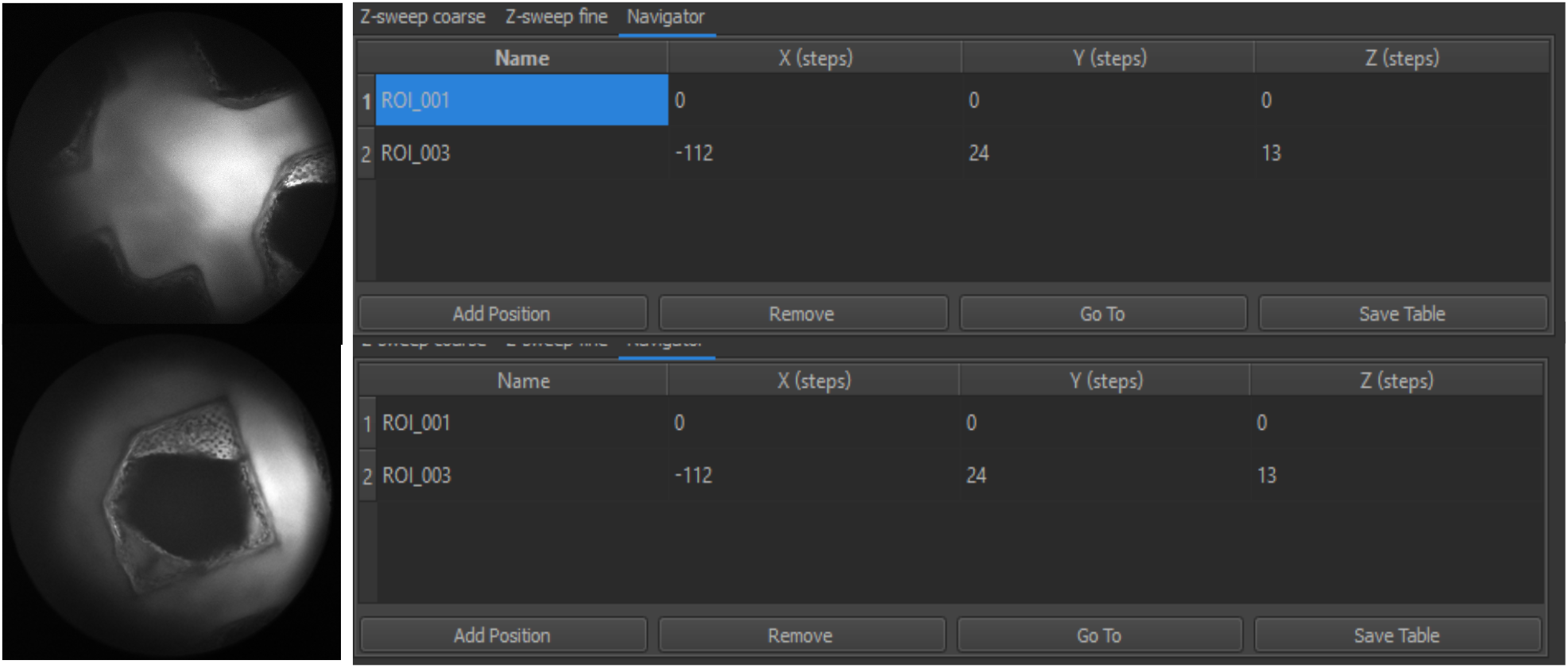
Navigation tab. The upper panel shows position 1 (‘ROI_001’), corresponding to the EM grid center at (x = 0, y = 0, z = 0). The lower panel shows position 2 (‘ROI_003’), corresponding to an EM grid square. The displayed x, y, and z values represent the number of steps at a set voltage of 20 V, which correspond to displacements of (−22.4, 4.8, 2.6) μm.

